# Alternative isoforms of KDM2A and KDM2B lysine demethylases negatively regulate canonical Wnt signaling

**DOI:** 10.1101/2020.07.13.200360

**Authors:** Dijana Lađinović, Daniel Pinkas, Otakar Raška, František Liška, Ivan Raška, Tomáš Vacík

**Affiliations:** Institute of Biology and Medical Genetics, First Faculty of Medicine, Charles University and General University Hospital in Prague, Albertov 4, Praha 2, Czech Republic

## Abstract

A precisely balanced activity of canonical Wnt signaling is essential for a number of biological processes and its perturbation leads to developmental defects or diseases. Here, we demonstrate that alternative isoforms of the KDM2A and KDM2B lysine demethylases have the ability to negatively regulate canonical Wnt signaling. These KDM2A and KDM2B isoforms (KDM2A-SF and KDM2B-SF) lack the N-terminal demethylase domain, but they are able to bind to activated promoters in order to repress them. We have observed that KDM2A-SF and KDM2B-SF bind to and repress the promoters of AXIN2 and CYCLIN D1, two canonical Wnt signaling target genes. Moreover, KDM2A-SF and KDM2B-SF can repress a Wnt-responsive luciferase reporter. The transcriptional repression mediated by KDM2A-SF and KDM2B-SF, but also by KDM2A-LF, is dependent on their DNA binding domain, while the N-terminal demethylase domain is dispensable for this process. Surprisingly, KDM2B-LF is unable to repress both the endogenous promoters and the luciferase reporter. Finally, we show that both KDM2A-SF and KDM2B-SF are able to interact with TCF7L1, one of the transcriptional mediators of canonical Wnt signaling. KDM2A-SF and KDM2B-SF are thus likely to affect the transcription of the TCF7L1 target genes also through this interaction.

## Introduction

KDM2A and KDM2B (KDM2A/B) are two closely related lysine demethylases with the ability to bind to non-methylated CpG islands through their CXXC DNA binding domain. After binding to non-methylated CpG islands in transcriptionally active promoters, KDM2A/B demethylate mono- and di-methylated H3K36 lysines (H3K36me1/2) using their N-terminal Jumonji-C demethylase domain (1, 2). KDM2B is able to demethylate also H3K4me3 (3). By demethylating H3K36me1/2 and H3K4me3 in transcriptionally active promoters (4–6), KDM2A/B function as transcriptional repressors of the promoters that contain CpG islands (1, 7–10). Although KDM2A and KDM2B have very similar structure, they have been shown to interact with different protein partners to repress different target regions. For example, KDM2A interacts with HP1a to repress the pericentromeric heterochromatin (11–13), whereas KDM2B forms complex with the PRC1 complex to silence developmentally important genes in embryonic stem cells (10).

Interestingly, KDM2A has been shown to interact with and to demethylate also non-histone proteins such as the p65 subunit of NF-kappaB or beta-catenin (14, 15). Beta-catenin is the key mediator of canonical Wnt signaling, which plays an essential role in a number of processes ranging from embryogenesis to aging, and whose malfunction frequently leads to various developmental defects and diseases including cancer (16–21). After activation of the pathway by Wnt ligands, beta-catenin enters the nucleus where it teams up with TCF/LEF transcription factors (TCF7L1, TCF7L2, TCF7, and LEF1) to activate transcription of their target genes. In the absence of Wnt ligands beta-catenin is phosphorylated at its N-terminal serines and threonines by the GSK-3/CKI kinases, which subsequently leads to its ubiquitination and finally to its proteasome mediated degradation. In the absence of beta-catenin in the nucleus, TCF/LEF proteins team up with co-repressors to act as transcriptional repressors of their target genes (16, 19, 22, 23). Interestingly, it has been demonstrated that KDM2A can displace the nuclear beta-catenin from the complex with TCF7L1, which results in transcriptional repression of the TCF7L1 target genes (14).

The same loci that encode the full-length KDM2A/B proteins (KDM2A/B-LF) also encode shorter KDM2A proteins that lack the N-terminal demethylase domain (1). However, these alternative short isoforms (KDM2A/B-SF) share all the other functional domains with KDM2A/B-LF. Therefore, KDM2A/B-SF are not able to demethylate the KDM2A/B target lysines, but they still have the ability to bind to the same DNA regions as KDM2A/B-LF and to interact with the same proteins as KDM2A/B-LF. KDM2A/B-SF are thus likely to compete with KDM2A/B-LF or to complement their function (1). Despite the fact that a KDM2A-SF specific knockout mutant has not been described yet and it cannot be compared to the embryonically lethal KDM2A-LF knockout phenotype (24), we previously showed that the distinct KDM2A positive nuclear structures on pericentromeric heterochromatin are formed by KDM2A-SF and not by KDM2A-LF (12). Similarly, the fact that the KDM2B-SF specific knockout phenotype is different from that of the KDM2B-LF loss-of-function mutants also implies different functions for the short and long isoforms (25–27).

Here we asked whether KDM2A/B-SF also affect canonical Wnt signaling despite lacking the demethylase domain. We demonstrate that both KDM2A-SF and KDM2B-SF have the ability to negatively regulate this signaling pathway by binding to and repressing the CpG island containing promoters of the pathway components AXIN2 and CYCLIN D1. Moreover, we found that KDM2A-SF and KDM2B-SF can interact with TCF7L1, one of the TCF/LEF transcriptional mediators of the pathway, which further broadens the negative effects that KDM2A/B-SF have on the TCF/LEF target genes.

## Material and Methods

### Cells

HEK293T cells were grown in 5% CO2 at 37°C and in the high glucose DMEM medium supplemented with 10% fetal bovine serum and the PenStrep antibiotics (all ThermoFisher Scientific). To stabilize beta-catenin and to induce canonical Wnt signaling the cells were treated with 1 μM BIO (6-Bromoindirubin-3′-oxime, SIGMA B1686) for 24 hours before harvesting.

### Plasmids and transfection

The coding regions of the corresponding genes were amplified by RT-PCR using the primers listed in supplementary table S1. The RT-PCR products were cloned in the pCS2 expression plasmids and the constructs were verified by sequencing. The mutant constructs were prepared by PCR mutagenesis using the Phusion polymerase (ThermoFisher Scientific). The TOP5 and FOP5 luciferase constructs were described previously (28). The *AXIN2* promoter luciferase constructs were prepared by PCR using the primers listed in supplementary table S1. The PCR products were cloned into the luciferase pNL1.1 vector (Promega) and verified by sequencing. The plasmids were transfected into cells using Fugene6 (Promega) or Turbofect (ThermoFisher Scientific).

### RNA and Q-RT-PCR

Total RNA was prepared with TRIzol (ThermoFisher Scientific) according to the manufacturer’s instructions and reverse transcribed with the LunaScript RT SuperMix kit (NEB). cDNA was analyzed by quantitative PCR using the CFX96 Touch Real-Time PCR Detection System (BIO-RAD), PowerUP SYBR Green mix (ThermoFisher Scientific), and the primers listed in supplementary table S1. The results were analyzed using the CFX Maestro software (BIO-RAD) and are presented as means ±SD of at least three independent experiments. The significance was determined using the student t-test.

### Luciferase reporter assay

Cells were co-transfected with the pNL1.1 nano luciferase reporter constructs, expression constructs and control firefly construct using Fugene6 (Promega). The reporter assays were performed using the NanoGlo Dual luciferase system (Promega) and the Infinite 200 luminometer (Tecan). To induce the transcription of the canonical Wnt signaling target genes, the cells were either treated with the pathway agonist BIO (1 μM, 24 hrs, SIGMA B1686) or transfected with the pCS2-beta-catenin expression constructs. The results are presented as means ±SD of at least three independent experiments.

### Proteins and western blot

Whole cell extracts were prepared by rotating the cell pellets for 2 hrs at + 4°C in five volumes of the high salt lysis buffer (50 mM Tris, 300 mM NaCl, 10% glycerol, 0.5% NP-40, 1x cOmplete ULTRA protease inhibitors (Roche)). Proteins were resolved on 10% SDS-PAGE gels, transferred to the Immobilon-P/E PVDF membrane (Merck Millipore), and immunodetected using the SuperSignal West Pico PLUS Chemiluminescent Substrate (ThermoFisher Scientific) and the following antibodies: anti-FLAG M2 (Sigma, F1804), anti-DYKDDDDK Tag (Cell Signaling, 14793), anti-Myc Tag (Millipore, 05-724), anti-mouse-HRP (GE healthcare, NA931-1ML), anti-rabbit-HRP (GE healthcare, NA934-1ML).

### Co-immunoprecipitation

HEK293T cells were transfected with the corresponding FLAG tag expression constructs using Turbofect (ThermoFisher Scientific). The whole cell extracts were prepared as described in 2.6. and diluted to 150 mM NaCl and 0.1% NP-40. 500 μg of the whole cell extract was rotated overnight at + 4°C with 2.5 μg of the anti-FLAG antibody (Sigma F1804) and the protein-immunocomplexes were separated with the Dynabeads Protein G magnetic beads (ThermoFisher Scientific). Proteins were eluted by boiling the beads in the LDS sample buffer (ThermoFisher Scientific) and analyzed by western blot.

### Chromatin immunoprecipitation

HEK293T cells were transfected with the pCS2-FLAG empty and protein coding constructs using Turbofect (ThermoFisher Scientific). After 48 hrs the cells were crosslinked with 1% formaldehyde (ThermoFisher Scientific) for 15 minutes at room temperature. The crosslinking reaction was stopped by 0.125M glycine and the samples were processed using the MAGnify Chromatin Immunoprecipitation System (ThermoFisher Scientific), Bioruptor (Diagenode), the anti-FLAG antibody (Sigma, F1804), anti-H3K4me3 (Cell Signaling, 9751), and the control IgG (Sigma, 12-371). The immunoprecipiated DNA was analyzed using the CFX96 Touch Real-Time PCR Detection System (BIO-RAD), Luna Universal qPCR Master Mix (NEB), and the primers listed in supplementary table S1. The HEK293T cells transfected on the same day with the same constructs were used to verify the expression of the FLAG-tagged proteins by western blot.

## Results

### KDM2A-SF and KDM2B-SF repress a Wnt-Responsive Luciferase Reporter

Since KDM2A-LF has been previously shown to strongly repress the Wnt-responsive luciferase Topflash reporter activated by elevated levels of beta-catenin (14), we set out to test whether the KDM2A-SF and KDM2B-SF isoforms are also able to repress this reporter despite lacking the demethylase domain. We used the TCF/LEF luciferase reporter TOP5, in which the luciferase gene is under the control of five TCF/LEF consensus binding sites and which thus reflects the activity of canonical Wnt signaling (Fig.1A)(28). We activated the canonical Wnt pathway in HEK293T cells with the pathway agonist BIO (6-Bromoindirubin-3′-oxime). BIO blocks the function of glycogen synthase kinase-3 (GSK3), whose role is to phosphorylate beta-catenin in the absence of a Wnt ligand and by doing so to prevent it from entering the nucleus (16, 18). Blocking the function of GSK3 results in accumulation of non-phosphorylated beta-catenin, its nuclear deposition and consequently in activation of TCF/LEF target genes including the above-mentioned TCF/LEF responsive TOP5 luciferase reporter (14, 29–34). Our luciferase reporter experiments confirmed that elevated levels of KDM2A-LF lead to a repression of the activated reporter, but they further showed that both KDM2A-SF or KDM2B-SF are also able to strongly repress this Wnt-responsive reporter despite lacking the N-terminal demethylase domain (Figs.1B and 1C). Interestingly, the full-length KDM2B-LF protein was not able to repress the activated TOP5 reporter (Fig.1C).

**Fig.1.**
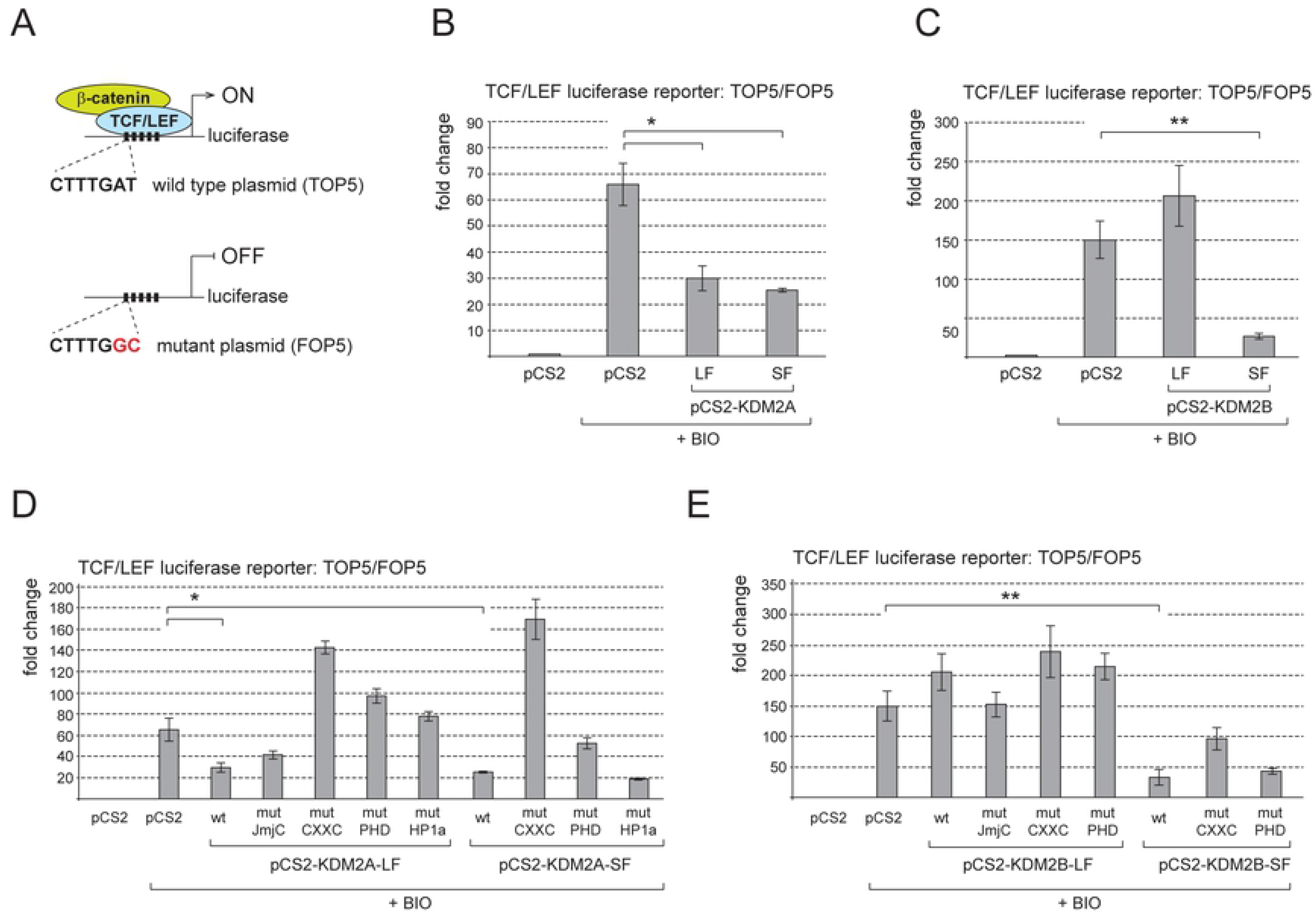
KDM2A-SF and KDM2B-SF repress the canonical Wnt signaling luciferase reporter. **A.** The wild type TCF/LEF reporter (TOP5) contains five TCF/LEF consensus binding sites (CTTTGAT) that drive the expression of the luciferase gene. The FOP5 construct contains five mutant TCF/LEF binding sites (CTTTGCC) instead and serves as the background activity control. **B.** Both KDM2A-LF and KDM2A-SF strongly repressed the TOP5 reporter activated with BIO. The reporter activity is expressed as the fold change ratio between the normalized luciferase signal of TOP5 and FOP5. **C.** KDM2B-SF, but not KDM2B-LF, also strongly repressed the activated reporter. **D.** Repression of the TOP5 reporter by KDM2A is not dependent on the activity of its JmjC demethylase domain, but predominantly on the CXXC DNA binding domain. **E.** KDM2B was not able to repress the activated TOP5 reporter, whereas KDM2B-SF strongly repressed it in a DNA binding domain dependent manner. (*p < 0.05; **p < 0.01).

We observed similar repression patterns when the TOP5 reporter was stimulated by the overexpression of beta-catenin (data not shown). These results imply that the N-terminal demethylase domain of KDM2A is not behind the transcriptional repressive effect that KDM2A has on the TOP5 reporter. To analyze that the activity of the JmjC demethylase domain is not required for the repressive effect, we performed the same reporter experiment with the KDM2A protein bearing the mutations previously shown to disrupt the function of the KDM2A demethylase domain, H212A and D214A (35). This experiment confirmed that the activity of the JmjC demethylase domain is dispensable for the KDM2A-mediated repression of the TOP5 reporter (Fig.1D). Surprisingly, the K601A mutation that disrupts the CXXC DNA binding domain of KDM2A not only reverted the repressive effect of both KDM2A-LF and KDM2A-SF, but it had a positive regulatory effect on the TOP5 reporter (Fig.1D, mutJmjC)(35). Similarly, disruption of the PHD domain by the C620A/C623A mutations abolished the repressive abilities of both KDM2A-LF and KDM2A-SF (Fig.1D, mutPHD)(11). Unlike KDM2B, KDM2A contains a short aminoacid motif that is important for the interaction with the heterochromatin protein HP1 (11). Disruption of this HP1 motif by the V801A and V803A substitutions reverted the repressive effect of KDM2A-LF, but not that of KDM2A-SF (Fig.1D, mutHP1a). As already stated above, KDM2B-LF did not repress the TOP5 reporter and mutating its functional domains had no significant effect in this regard (Fig.1E). On the other hand, the strong repression of the TOP5 reporter by KDM2B-SF was also dependent on its DNA-binding domain, whereas its PHD domain seems dispensable for this repression (Fig.1E).

### KDM2A-SF and KDM2B-SF repress transcription of AXIN2 and CYCLIN D1

To complement our luciferase reporter data and to analyze whether KDM2A-SF and KDM2B-SF are able to repress also endogenous canonical Wnt signaling target genes, we focused on AXIN2 and CYCLIN D1. AXIN2 and CYCLIN D1 are two widely studied direct target genes of canonical Wnt signaling and as such their promoters can be activated with BIO (36–40). We stimulated their expression with BIO and analyzed their transcriptional activity by Q-RT-PCR in the presence of the above described KDM2A and KDM2B protein variants. Consistently with our luciferase assays, both KDM2A-LF and KDM2A-SF strongly repressed activated AXIN2 and CYCLIN D1 (Figs.2A and 2B). While the transcriptional repression of AXIN2 by both KDM2A-LF and KDM2A-SF is dependent on their CXXC DNA binding domain, disruption of this domain did not completely revert the repressive effect on CYCLIN D1 (Figs.2A and 2B, mutCXXC). However, the KDM2A-LF mediated repression of the *CYCLIN D1* gene seems to be dependent on the HP1 motif, whose disruption reverted the repressive effect (Fig.2B, mutHP1a). This implies that the nature of the *CYCLIN D1* transcriptional regulation by the KDM2A isoforms is different from that of AXIN2. Consistently with the results of our luciferase assay, KDM2B-LF failed to repress both AXIN2 and CYCLIN D1 (Figs.2C and 2D). On the other hand, KDM2B-SF efficiently repressed both AXIN2 and CYCLIN D1 in a DNA binding domain dependent manner, since disruption of its CXXC domain by the C586/589/592A mutations reverted the repression (Figs.2C and 2D)(41). The C661/664A mutations that disrupt the KDM2B PHD domain had no significant effect and these KDM2B mutant proteins still repressed both AXIN2 and CYCLIN D1 (Figs.2C and 2D)(42).

**Fig.2.**
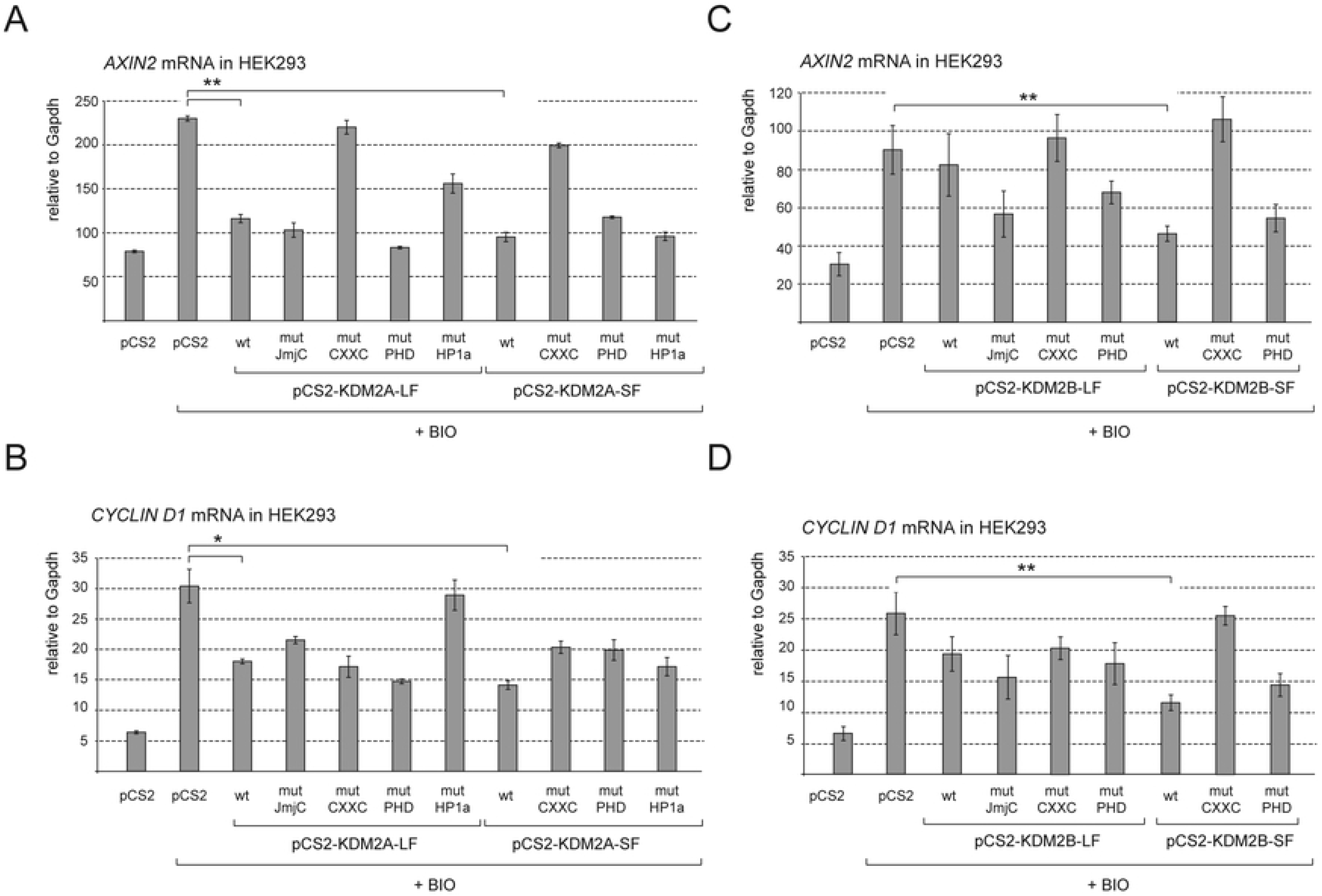
KDM2A-SF and KDM2B-SF repress AXIN2 and CYCLIN D1. **A.** Both KDM2A-LF and KDM2A-SF repressed AXIN2 in a DNA binding domain dependent manner. **B.** When activated with BIO, CYCLIN D1 is also repressed by both KDM2A-LF and KDM2A-SF, but independently of the DNA binding domain. **C.** Similarly to KDM2A-SF, KDM2B-SF repressed activated AXIN2, whereas KDM2B-LF had no effect on its transcription. **D.** CYCLIN D1 was also repressed by KDM2B-SF, but not by KDM2B-LF. The mRNA levels were determined by Q-RT-PCR and are related to GAPDH. Similar patterns were obtained also with HPRT and RPL32 (*p < 0.05; **p < 0.01).

### KDM2A-SF and KDM2B-SF repress AXIN2 promoter

Transcriptional repression of AXIN2 and CYCLIN D1 by KDM2A-LF, KDM2A-SF and KDM2B-SF is well consistent with the fact that KDM2A/B act as CpG island binding transcriptional repressors (1). Since the *AXIN2* and *CYCLIN D1* promoters contain CpG islands, the KDM2A/B-SF mediated repression of these promoters is likely to be direct. To investigate whether KDM2A-SF and KDM2B-SF directly repress the *AXIN2* promoter, we focused on the *AXIN2* promoter region in its first intron (Fig.3A). This intronic promoter region contains a CpG island and it has been shown to act as an important regulatory promoter element of AXIN2 (38). The ENCODE ChIP-seq data that are publicly available via the UCSC genome browser further show that this region is bound by RNA-POL II and various transcription factors in multiple cell lines, which further confirms its importance for the transcriptional regulation of AXIN2 (43). Consistently with the above mentioned facts, this approximately 2.5 kb *AXIN2* intron 1 region exhibited a high transcription inducing activity in a luciferase assay as opposed to a shorter approximately 1 kb *AXIN2* intron 1 region that lacks the CpG island (Figs.3A and 3B). Consistently with our TOP5 reporter and Q-RT-PCR results, the luciferase activity driven by this *AXIN2* promoter region was repressed by KDM2A-LF, KDM2A-SF, and KDM2B-SF, whereas KDM2B-LF was not able to repress this region (Fig.3B).

**Fig.3.**
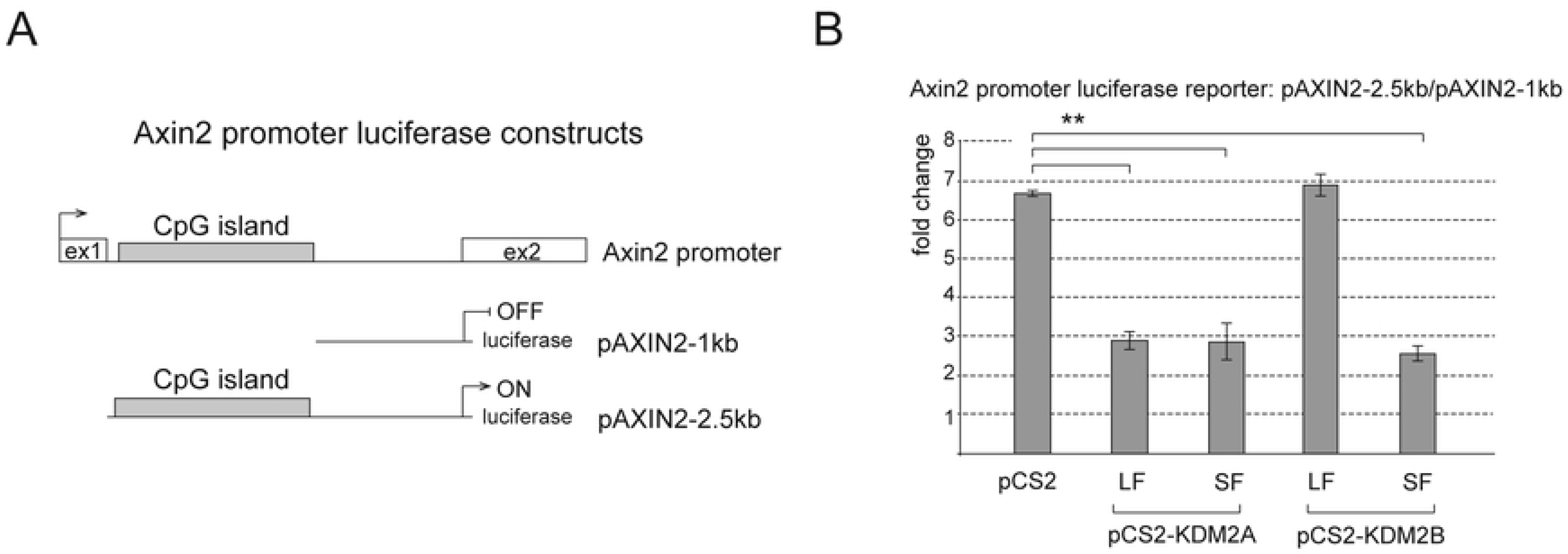
KDM2A-SF and KDM2B-SF repress AXIN2 promoter. **A.** The *AXIN2* promoter luciferase reporter constructs contain either a 1kb *AXIN2* intron 1 region (pAXIN2-1kb) or a 2.5kb *AXIN2* intron 1 region (pAXIN2-2.5kb). **B.** KDM2A-LF, KDM2A-SF and KDM2B-SF all repressed the pAXIN2-2.5kb reporter, whereas KDM2B-LF did not. The activity of the pAXIN2-2.5kb reporter is expressed as the fold change ratio between the activity of pAXIN2-2.5kb and that of pAXIN2-1kb. Similar pattern was observed when the results were expressed as the fold change between the activity of pAXIN2-2.5kb and the empty luciferase plasmid. (*p < 0.05; **p < 0.01).

### KDM2A-SF and KDM2B-SF bind to the repressed AXIN2 and CYCLIND1 promoter regions

Our results imply that KDM2A-SF and KDM2B-SF directly repress the transcription of AXIN2 and CYCLIN D1. To test whether KDM2A/B-SF isoforms directly bind to the promoters of these genes we performed a series of chromatin immunoprecipitation (ChIP) assays. The amino acid sequence of the alternative KDM2A/B-SF isoforms is identical to that of the corresponding region of the canonical full-length KDM2A/B-LF proteins (1). Therefore, it is not possible to prepare antibodies specific for KDM2A-SF and KDM2B-SF. To discriminate between binding of the long and short isoforms of KDM2A/B, we overexpressed their N-terminally FLAG-tagged versions in HEK293T cells and tested the selected regions for their presence by ChIP. In our ChIP assay we focused on the promoter regions that have been previously shown to be important for transcriptional regulation of AXIN2 and CYCLIN D1 (38–40). Furthermore, our *in silico* analysis of the publicly available data showed that these regions contain CpG islands, which makes them potential targets of the KDM2A/B CpG island binding proteins, and that they are bound by multiple transcription factors in various cell lines, which further confirms their role in transcriptional regulation of the associated genes (43). Consistently with the results of our Q-RT-PCR and luciferase experiments, the ChIP assay showed that KDM2A-LF, KDM2A-SF and KDM2B-SF bind to the promoter of AXIN2, whereas KDM2B-LF does not (Figs.4A and 4B). Moreover, we tested the same *AXIN2* promoter region for the levels of H3K4me3, the histone lysine methylation associated with transcriptionally active regions (4, 5). This ChIP experiment revealed that the H3K4me3 levels on the *AXIN2* promoter expectedly rise after the treatment with BIO, whereas they fall back in the presence of KDM2A-LF or KDM2A-SF (Fig.4C). These changes in the H3K4me3 levels are consistent with the transcriptional activation of the *AXIN2* promoter with BIO and with the KDM2A/B-SF mediated repression of this promoter, respectively. Similarly, the *CYCLIN D1* promoter was bound by KDM2A-LF, KDM2A-SF and KDM2B-SF, but not by KDM2B-LF (Figs.4D and 4E), and its H3K4me3 levels are also lower in the presence of KDM2A-LF or KDM2A-SF (Fig.4F).

**Fig.4.**
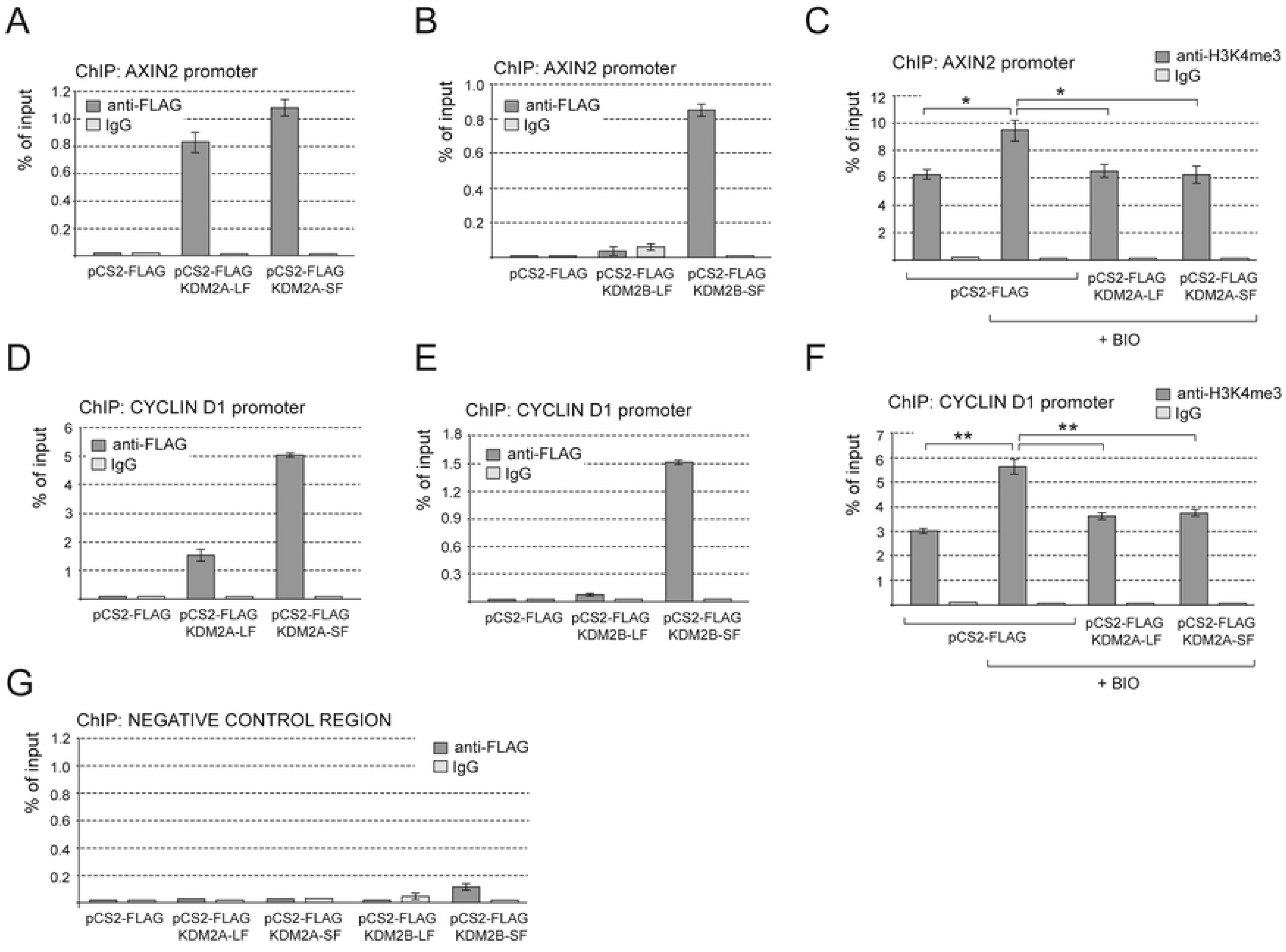
KDM2A-SF and KDM2B-SF bind to the repressed AXIN2 and CYCLIND1 promoter regions. **A.** KDM2A-LF and KDM2A-SF both bind to the *AXIN2* promoter. **B.** Only KDM2B-SF, but not KDM2B-LF, binds to the *AXIN2* promoter. **C.** The levels of H3K4me3 on the *AXIN2* promoter rose after the treatment with BIO and then decreased in the presence of the KDM2A isoforms. **D.** KDM2A-SF binds to the *AXIN2* promoter with a higher efficiency than KDM2A-LF. **E.** KDM2B-SF, but not KDM2B-LF, binds to the *CYCLIN D1* promoter. **F.** The KDM2A isoforms negatively affected the H3K4me3 levels also on the *CYCLIN D1* promoter. **G.** The negative control region did not co-immunoprecipitate with the KDM2A/B isoforms. (*p < 0.05; **p < 0.01).

### KDM2A-SF and KDM2B-SF interact with TCF7L1

Since KDM2A-LF has been shown to form a complex with TCF7L1 (14), we set out to investigate whether KDM2A-SF and KDM2B-SF are also able to interact with this transcriptional mediator of canonical Wnt signaling. Our co-immunoprecipitation (Co-IP) assays confirmed that tagged KDM2A-LF and TCF7L1 overexpressed in HEK293T interact (Fig.5). In addition, our Co-IP experiments revealed that both short isoforms, KDM2A-SF and KDM2B-SF, and the canonical full-length KDM2B-LF isoform are also able to form complex with TCF7L1 (Fig.5). These results imply that the N-terminal demethylase domain of the KDM2A/B demethylases is not necessary for the interaction with TCF7L1. Furthermore, these interactions help explain why KDM2A-SF and KDM2B-SF can repress the TCF/LEF-responsive TOP5 luciferase reporter (Fig.1), although the promoter of this reporter does not contain any CpG island.

**Fig.5.**
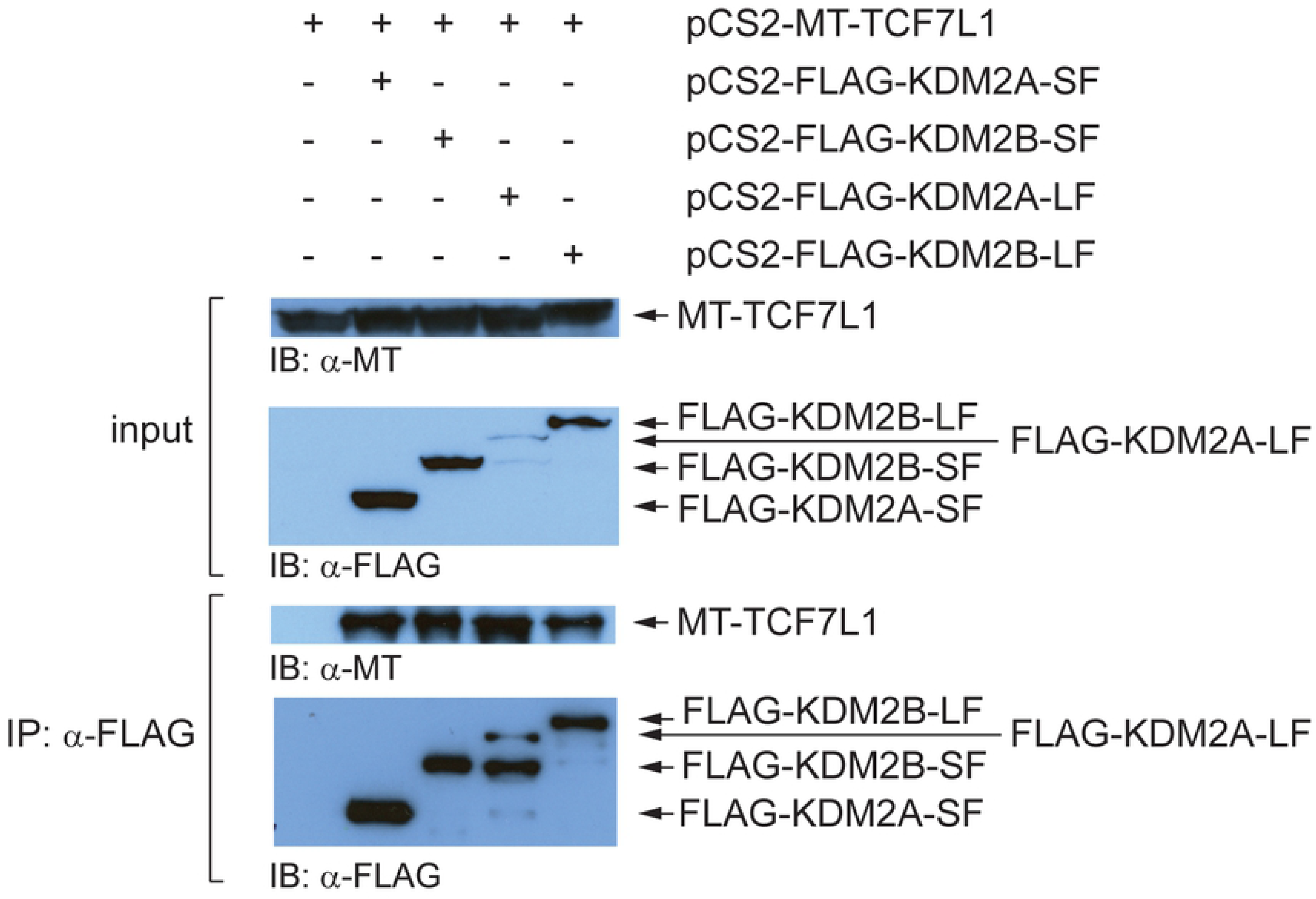
Both the canonical and the demethylase domain deficient isoforms of KDM2A and KDM2B interact with TCF7L1. The myc-tagged TCF7L1 was overexpressed in HEK293T cells together with the FLAG-tagged KDM2A/B isoforms and the proteins immunoprecipitated with anti-FLAG antibody were analyzed by western blot. All the four FLAG-tagged KDM2A/B isoforms co-immunoprecipitated with TCF7L1. The input corresponds to 5% of the whole cell extract that was used for each immunoprecipitation reaction.

## Discussion

In this study, we demonstrate that KDM2A-SF and KDM2B-SF, the two alternative isoforms of the lysine demethylases KDM2A and KDM2B that lack the demethylase domain, are able to negatively affect canonical Wnt signaling at the transcriptional level. We demonstrate that KDM2A-SF and KDM2B-SF bind to and repress the promoters of the canonical Wnt signaling target genes AXIN2 and CYCLIN D1. The *AXIN2* and *CYCLIN D1* promoter regions we focused on here have been previously studied and shown to be important for transcriptional regulation of the two genes (38–40). The regulatory function of these regions is further supported by the publicly available ENCODE ChIP-seq data, which indicate that these regions are bound by RNA-POL II and multiple transcription factors in various cell lines (43). Moreover, we found that these regulatory regions contain CpG islands, which makes them potential targets of KDM2A and KDM2B, and also of KDM2A-SF and KDM2B-SF, since all these protein isoforms are CpG island binding proteins. The binding of KDM2A and KDM2B to these CpG islands in mouse embryonic stem cells have already been experimentally verified by ChIP-seq (10, 35). However, pan-antibodies that recognize both the canonical long (KDM2A/B-LF) and the short (KDM2A/B-SF) isoforms were used in these studies and they are thus not informative as to what isoforms are bound to these promoter regions.

Since KDM2A-SF and KDM2B-SF have the identical aminoacid sequence as the corresponding regions of the long KDM2A/B-LF isoforms (1), it is not possible to prepare antibodies specific just for KDM2A/B-SF. To circumvent this obstacle, we used FLAG-tagged versions of KDM2A/B-SF and KDM2A/B-LF proteins to immunoprecipitate the chromatin regions bound by them by ChIP. Our ChIP assay demonstrated that both isoforms of KDM2A, KDM2A-LF and KDM2A-SF, bind to the CpG island containing promoter region of both AXIN2 and CYCLIN D1 (Figs.4A and 4D), which is consistent with the ability of these protein isoforms to repress the transcription driven by these promoters (Fig.2)., and to repress the *AXIN2* promoter luciferase construct (Fig3). Using ChIP, we also analyzed the selected regions for the levels of H3K4me3, a mark of transcriptionally active promoter regions (4, 5). Consistently with the transcriptional activation of AXIN2 and CYCLIN D1 upon stimulation of canonical Wnt signaling (Fig.2), the H3K4me3 levels in the tested promoter regions rose after the treatment with BIO (Fig.4). The ChIP assay showed that the presence of either KDM2A-LF or KDM2A-SF results in statistically lower H3K4me3 levels, which corresponds to the transcriptionally repressive effect of these proteins on the tested promoters (Fig.4). We detected similar results also for KDM2B-SF, whose binding to the two tested promoter regions leads to lower H3K4me3 levels (data not shown) and to transcriptional repression of the corresponding genes (Fig.2). Surprisingly, KDM2B-LF was unable to bind to the tested promoter regions, which is consistent with the fact that this canonical long KDM2B protein isoform failed to repress the transcription of AXIN2 and CYCLIN D1 (Fig.2). The direct repressive effect of KDM2A-LF, KDM2A-SF, and KDM2B-SF on the *AXIN2* promoter is further supported by our luciferase assays, which show that the *AXIN2* promoter region is repressed by these protein isoforms, whereas KDM2B-LF again failed to show any transcriptionally repressive properties (Fig.3).

These results are further consistent with the fact that KDM2A-LF, KDM2A-SF, and KDM2B-SF, but not KDM2B-LF, are able to repress the stimulated TOP5 luciferase reporter (Fig.1). However, the TOP5 reporter does not contain any CpG island and so the repressive effect is either not dependent on the CpG island binding domain or it is mediated by some auxiliary protein. Our luciferase data demonstrate that KDM2A-SF and KDM2B-SF need their DNA binding domain to repress the reporter and the KDM2A PHD domain is also involved to some extent (Fig.1). Based on the previously described interaction of KDM2A-LF with TCF7L1 (14), and on the fact that the luciferase gene is driven from the TOP5 plasmid by five TCF/LEF binding sites, we hypothesized that the repression of the TOP5 reporter by KDM2A/B-SF might be mediated by TCF7L1. Therefore, we tested whether KDM2A/B-SF are also able to interact with TCF7L1 by Co-IP. Our Co-IP results indeed show that all the four KDM2A/B isoforms can interact with TCF7L1 (Fig.5). KDM2A/B-SF are thus likely to repress the TCF/LEF reporter, but also endogenous TCF/LEF target promoters, by interacting with TCF7L1. Our Co-IP results further indicate that the N-terminal region of KDM2A/B is not necessary for the interaction with TCF7L1.

In this study, we present a mechanism that regulates the canonical Wnt signaling activity at the transcriptional level through KDM2A-SF and KDM2B-SF. KDM2A-SF and KDM2B-SF have the ability to affect canonical Wnt signaling by both binding to the promoters of canonical Wnt signaling target genes such as AXIN2 or CYCLIN D1, and by interacting with TCF7L1.

